# Single synapses control mossy cell firing

**DOI:** 10.1101/188789

**Authors:** Alexander Drakew, Urban Maier, Anja Tippmann, Michael Frotscher

**Author notes:** both authors contributed equally to this study (Alexander Drakew, Urban Maier). Correspondence to: A.D. Michael Frotscher passed away on May, 27 2017.

## Abstract

The function of mossy cells (MCs) in the dentate gyrus has remained elusive. Here we determined the functional impact of single mossy fibre (MF) synapses on MC firing in mouse entorhino-hippocampal slice cultures. We stimulated single MF boutons and recorded Ca^2+^ transients in the postsynaptic spine and unitary excitatory postsynaptic potentials (EPSPs) at the MC soma. Synaptic responses to single presynaptic stimuli varied strongly between different MF synapses, even if they were located on the same MC dendrite. Synaptic strengths ranged from subthreshold EPSPs to direct postsynaptic action potential (AP) generation. Induction of synaptic plasticity at these individual MF synapses resulted in potentiation or depression depending on the initially encountered synaptic state, indicating that synaptic transmission at MF synapses on MCs is determined by their previous functional history. With these unique functional properties MF-MC synapses control MC firing individually thereby enabling modulation of the dentate network by single granule cells.

---

Sensory input that reaches the entorhinal cortex from visual, olfactory and auditory centres is relayed to the hippocampus via distinct connectivity channels formed by the synapses of dentate granule cells^1^. The mossy fibres (MFs), the axons of the granule cells, give rise to giant presynaptic boutons that terminate on large complex spines on the proximal dendritic segments of pyramidal neurons^2–5^ and on dendritic shafts of GABAergic interneurons^6,7^ in the CA3 region. However, on their way to CA3 MFs first encounter the mossy cells (MCs) in the hilus of the dentate gyrus^8^. In contrast to CA3 pyramidal neurons, MCs feature MF synapses on all dendritic segments and on various types of spines^9^. MF-MC synapses are thought to be equivalent to MF-CA3 synapses in showing presynaptic expression of LTP and frequency facilitation^10,11^. Being embedded in the hilus and exposed to abundant bypassing MFs, single MCs may receive input from several hundred (800) granule cells^12^. Thus, the specific segregated granule cell output, which conveys information about contextual changes in the external environment from the entorhinal cortex to the hippocampus, converges on MCs^13–16^.

MCs are excitatory glutamatergic neurons^17^ that project to the hilus locally and to the inner molecular layer of distant portions of the ipsilateral and contralateral dentate gyrus^18,19^, innervating numerous granule cells and dentate inhibitory interneurons in both hemispheres. MCs are thought to modulate the excitation/inhibition balance of dentate granule cells differentially in local and distant parts of the dentate gyrus thereby promoting the formation of functional transversal lamellae in the hippocampus^8,20,21^. Consistent with this idea, genetic ablation of MCs resulted in transient granule cell hyperexcitability and impaired pattern separation^22^. Furthermore, MCs are the first excitatory neurons that activate newly generated granule cells, which extend their yet short dendrites to the inner molecular layer^23^. Comparative studies revealed no principal differences between MCs in rodents and primates^9^, suggesting that MCs are an important, conserved component of the hippocampal network.

A remarkable feature of MF synapses is their structural plasticity. Structural complexity of MF boutons was found to increase with the age of the animal and in response to enriched environment^24^. Similarly, in hippocampal slice cultures the complexity of MF boutons increased with the incubation period *in vitro* and with functional activity, thus establishing slice cultures as a tool to study MF synaptic plasticity^24,25^. In line with these results, stimulation of single granule cells increased the complexity of their respective MF boutons^26^. Earlier studies had already shown that the number and size of complex spines postsynaptic to MF boutons were reduced following removal of the entorhinal cortex^27^. Collectively these results suggest that MF synapses change their structural components individually depending on the activity of the parent granule cell in response to environmental stimuli mediated by the entorhinal cortex.

While these morphological studies revealed structural variability of MF synapses, they could not clarify whether individual MF synapses were functionally different. Functional synaptic modification, such as potentiation of synaptic transmission^28^ is assumed to underlie adaptive mechanisms such as learning and memory^29,30^ and is likely to occur at individual synapses depending on their previous individual involvement in active microcircuits. Hence, at a given point in time one would expect to envisage different, non-random states of synaptic transmission in a defined population of synapses, such as MF synapses. Moreover, little is known about the detonation properties, the efficacy, and plasticity of individual MF-MC synapses.

In the present study, we hypothesized that the synaptic strengths of individual MF synapses on MCs are heterogeneous and are modified individually by activity. We tested the functional behaviour of individual synapses by activating single MF boutons on MC dendrites, recording Ca^2+^ transients in spines postsynaptic to these boutons and unitary EPSPs at the MC soma, and inducing plasticity at these synapses using a defined stimulation protocol.

## Results

### Determining the synaptic states of individual MF synapses on MCs

We chose organotypic entorhino-hippocampal slice cultures as a model system to study the individual states of single synapses under stable *in vitro* conditions. Acute slices would have been an alternative model for studies of that kind. However, slice preparation has been shown to change acutely the number of spines and synapses in response to acute deafferentation by the transection of many axons and dendrites^31^. Thus, states of synaptic transmission may not reflect previous synaptic activity under these conditions. We used organotypic entorhino-hippocampal slice cultures that were incubated *in vitro* for at least 4 weeks to allow for recovery and stabilization of the tissue following slice preparation, the removal of debris, and for the maturation of the MF projection. Such long-term slice cultures were shown before to represent a reliable model for the study of MF synapses since the development and activity-induced structural plasticity of MF synapses were very similar *in vivo* and in slice culture^24^. In the present study we used hippocampal slice cultures with the entorhinal cortex attached to preserve major afferent projections to the granule cells which retain their layer-specific terminations under these *in vitro* conditions^32^. Moreover, the fine-structural characteristics of MF synapses were preserved in slice culture and there were no principal differences to MF synapses *in vivo*^33^. Accordingly, this *in vitro* model has been successfully used in studies on the functional and structural plasticity of MF synapses^24,34^.

MCs were patch-clamped and filled with Alexa 594 dextran for cell identification (Fig. 1a, d) and with Fluo-4FF for two-photon Ca^2+^ imaging (Fig. 1b, see Methods). Targeted patch-clamp recording of MF boutons contacting the labelled spines of the MC dendrite was facilitated by transiently staining the extracellular space with Alexa 488 hydrazide, which was not taken up by cellular elements, allowing to visualise the boutons as “shadows” surrounding the spines^35^ (Fig. 1b, c). Stimulation of MF boutons in loose-seal cell-attached mode was combined with recording of unitary EPSPs from the target MC soma and simultaneous two-photon imaging of Ca^2+^ transients in the postsynaptic spine (Fig. 1e, f, Supplementary Fig. 1a-c, Supplementary Table 1). EPSPs following bouton stimulation were indistinguishable from EPSPs in response to spontaneous input to the same cell (Supplementary Fig. 2a). Ca^2+^ transients observed in the imaged spines reliably reported synaptic events occurring at this particular synapse since suprathreshold EPSPs generated somewhere else in the dendritic tree elicited only very small Ca^2+^ transients in the spine when compared to local EPSPs (Supplementary Fig. 2b, c).

**Fig. 1.**
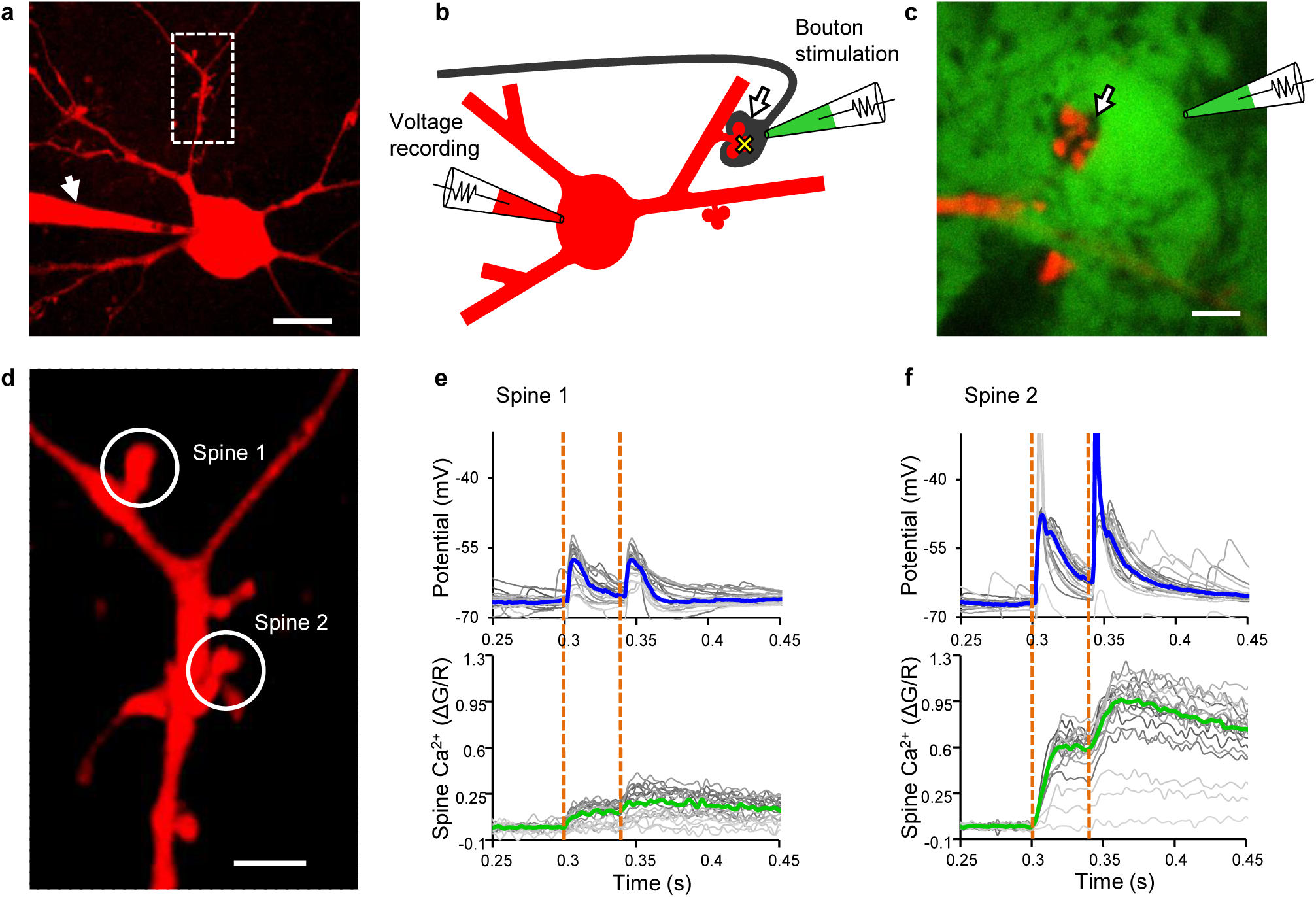
Interrogation of single MF-MC synapses. **a** MC filled with Alexa 594 dextran and the Ca^2+^-sensitive dye Fluo-4FF via a patch pipette (arrow). Scale bar: 20 μm. Boxed area shown in **d. b** Schematic diagram illustrating stimulation and recording sites. Arrow: MF bouton; yellow cross: imaging site of Ca^2+^ transients; green: extracellular tracer (Alexa 488 hydrazide). **c** Two-photon XY-frame to illustrate the bouton “shadow” (arrow) before patching. Scale bar: 5 μm. **d** Boxed area in **a**. Boutons presynaptic to the two encircled spines were stimulated and voltage responses recorded at the somatic pipette. Simultaneously, Ca^2+^ transients were recorded in the spines. Scale bar: 5 μm. **e**, **f** Unitary voltage responses and Ca^2+^ transients evoked by paired-pulse stimulation (orange dotted lines) of MF synapses on spine 1 and 2, respectively. Blue and green lines: Medians of all repetitions; grey lines: Individual trials.

**Table 1.**
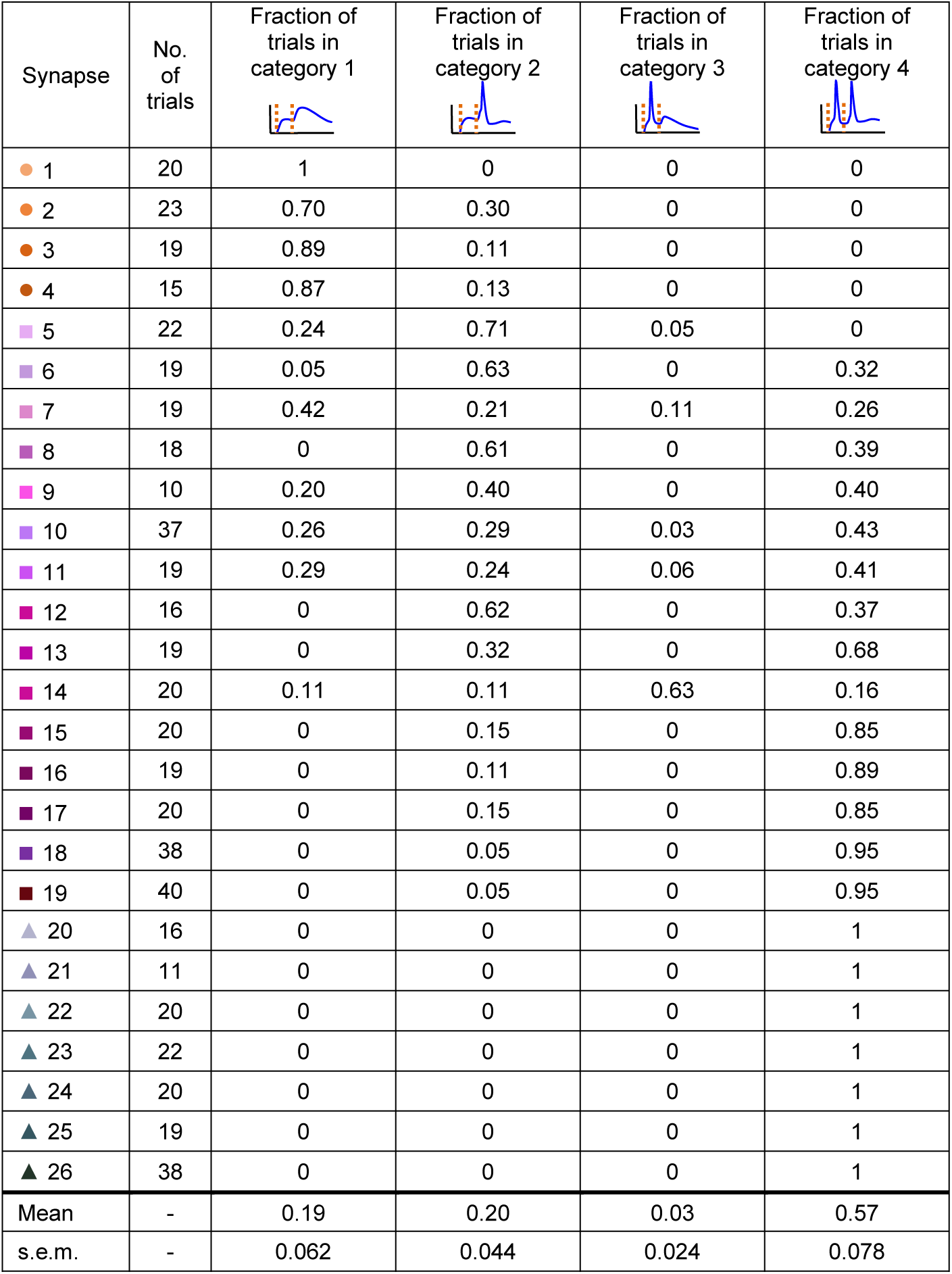
Classification of responses to paired-pulse stimulation. Response patterns are shown schematically on top (category 1: both responses are subthreshold; category 2: only the response to the second stimulus is suprathreshold; category 3: only the response to the first stimulus is suprathreshold; category 4: both responses are suprathreshold). Each synapse is represented in all 4 response categories (fractions summing up to 1). Coding of synapses by coloured symbols as in Fig. 2. Bottom: Mean ± s.e.m. of fractions of all 26 synapses.

| Synapse | No. of trials | Fraction of trials in category 1 | Fraction of trials in category 2 | Fraction of trials in category 3 | Fraction of trials in category 4 |
| --- | --- | --- | --- | --- | --- |
| ● 1 | 20 | 1 | 0 | 0 | 0 |
| ● 2 | 23 | 0.70 | 0.30 | 0 | 0 |
| ● 3 | 19 | 0.89 | 0.11 | 0 | 0 |
| ● 4 | 15 | 0.87 | 0.13 | 0 | 0 |
| ■ 5 | 22 | 0.24 | 0.71 | 0.05 | 0 |
| ■ 6 | 19 | 0.05 | 0.63 | 0 | 0.32 |
| ■ 7 | 19 | 0.42 | 0.21 | 0.11 | 0.26 |
| ■ 8 | 18 | 0 | 0.61 | 0 | 0.39 |
| ■ 9 | 10 | 0.20 | 0.40 | 0 | 0.40 |
| ■ 10 | 37 | 0.26 | 0.29 | 0.03 | 0.43 |
| ■ 11 | 19 | 0.29 | 0.24 | 0.06 | 0.41 |
| ■ 12 | 16 | 0 | 0.62 | 0 | 0.37 |
| ■ 13 | 19 | 0 | 0.32 | 0 | 0.68 |
| ■ 14 | 20 | 0.11 | 0.11 | 0.63 | 0.16 |
| ■ 15 | 20 | 0 | 0.15 | 0 | 0.85 |
| ■ 16 | 19 | 0 | 0.11 | 0 | 0.89 |
| ■ 17 | 20 | 0 | 0.15 | 0 | 0.85 |
| ■ 18 | 38 | 0 | 0.05 | 0 | 0.95 |
| ■ 19 | 40 | 0 | 0.05 | 0 | 0.95 |
| ▲ 20 | 16 | 0 | 0 | 0 | 1 |
| ▲ 21 | 11 | 0 | 0 | 0 | 1 |
| ▲ 22 | 20 | 0 | 0 | 0 | 1 |
| ▲ 23 | 22 | 0 | 0 | 0 | 1 |
| ▲ 24 | 20 | 0 | 0 | 0 | 1 |
| ▲ 25 | 19 | 0 | 0 | 0 | 1 |
| ▲ 26 | 38 | 0 | 0 | 0 | 1 |
| Mean | - | 0.19 | 0.20 | 0.03 | 0.57 |
| s.e.m. | - | 0.062 | 0.044 | 0.024 | 0.078 |

**Fig. 2.**
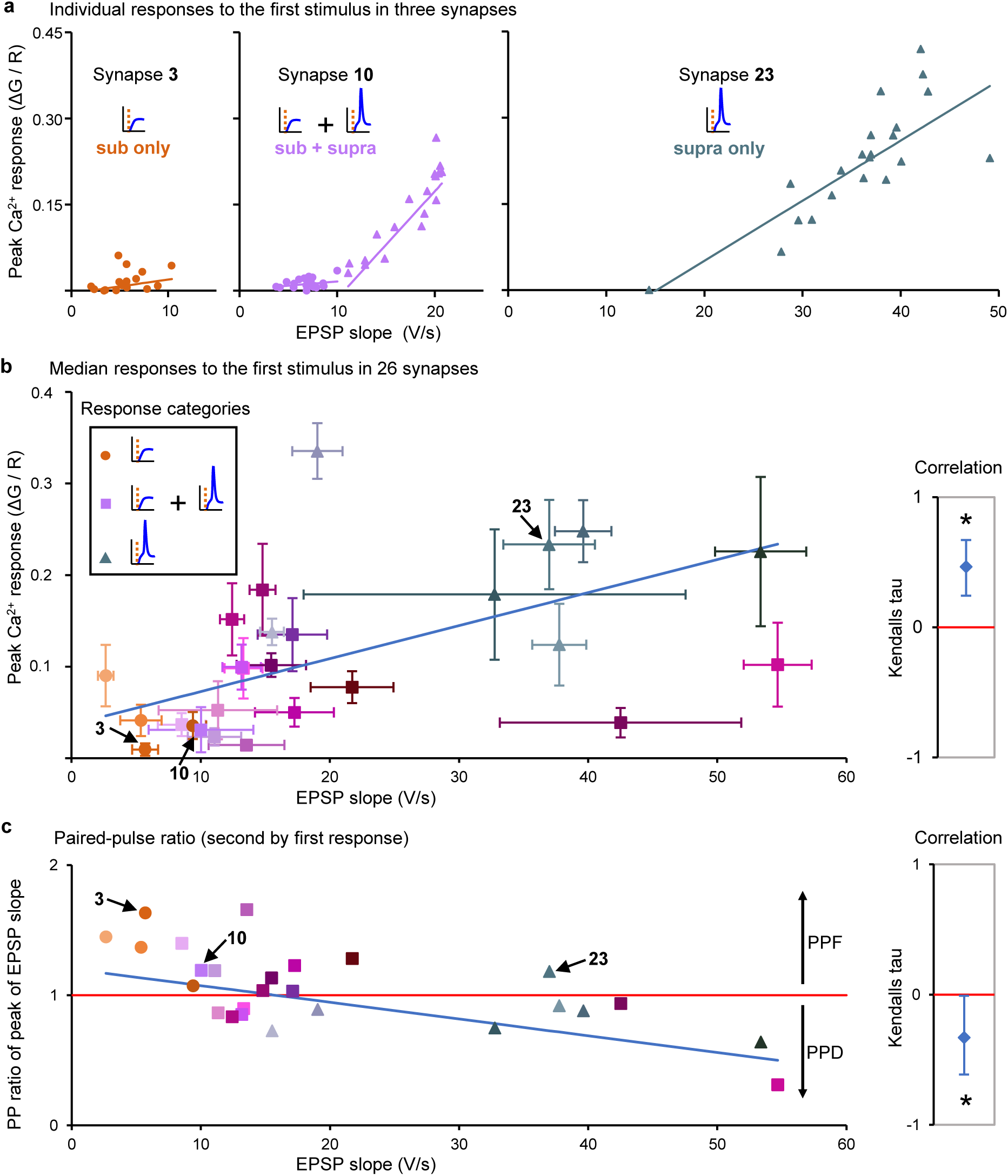
Functional heterogeneity of MF-MC synapses. **a** Unitary voltage and Ca^2+^ responses to the first of the paired stimuli in three MF-MC synapses from three different slice cultures (synapse 3, synapse 10, and synapse 23). (●) subthreshold responses, (▲) suprathreshold responses. Lines represent regression of peak Ca^2+^ amplitudes on EPSP slopes in the three synapses (separate regression lines for sub-and suprathreshold responses for synapse 10, showing supralinear Ca^2+^ rises if postsynaptic APs were generated). Sub: subthreshold response; supra: suprathreshold response. **b** Medians of all responses obtained in each of the 26 synapses from 26 different slice cultures, bars representing median absolute deviations from the median. The three synapses in **a** are indicated. Inset: (●) only subthreshold responses; (■) mixed responses; (▲) only suprathreshold responses to the first stimulus. Blue line represents regression of peak Ca^2+^ amplitude on EPSP slope, right panel shows the correlation between these parameters. The bootstrap mean (♦) of Kendalls tau and a confidence interval (0.025 – 0.975, vertical bars) is shown as a robust measure of correlation. If 0 (red line) is not included in the confidence interval, the parameters are considered to be correlated (*) at a significance level of 0.05. **c** Paired-pulse ratios (PPR) of EPSP slope versus EPSP slope of the 26 synapses (medians of all repetitions per synapse). Red line indicates a PPR of 1. Blue line represents regression of PPR of EPSP slope on EPSP slope. PPF and PPD: Paired-pulse facilitation and depression. Right panel shows the correlation between PPR of EPSP slope and EPSP slope of these synapses.

We used paired-pulse stimulation of the MF bouton to interrogate individual MF synapses since the response to the first of the two stimuli allowed us to capture the current state of the synapse, while the voltage response to the second stimulus reported presynaptic facilitation known to increase transmitter release probability at MF synapses^36^. Paired-pulse facilitation, expressed as ratio of the second to the first response (paired-pulse ratio), allowed us to probe the initial release probability^37^. Paired-pulse facilitation is a form of short-term synaptic plasticity that needs to be differentiated from long-term changes of synaptic strength. Accordingly, presynaptic facilitation did not accumulate with repetitive paired-pulse stimulations in the synapses we studied. However, the non-invasive loose-seal bouton-attached configuration turned out to be stable enabling us to probe transmission at the same synapse again following induction of long-term synaptic potentiation applying an associative protocol^38^. This approach allowed us to analyse simultaneously presynaptic and postsynaptic components of transmission before and after induction of synaptic plasticity at single identified synapses. Of note, these experiments were performed in ACSF without any pharmacological additives at near physiological temperature.

### MF synapses on MCs display heterogeneous synaptic states

MF synapses on CA3 pyramidal cells are “conditional detonators”, discharging the postsynaptic cell only in response to a train of granule cell APs^39–41^. In contrast, in 85% of the MF synapses on MCs we observed direct AP firing of the MC (“direct detonation”) in response to the first stimulus (Fig. 1f, Fig. 2a, b, Supplementary Fig. 3).

**Fig. 3.**
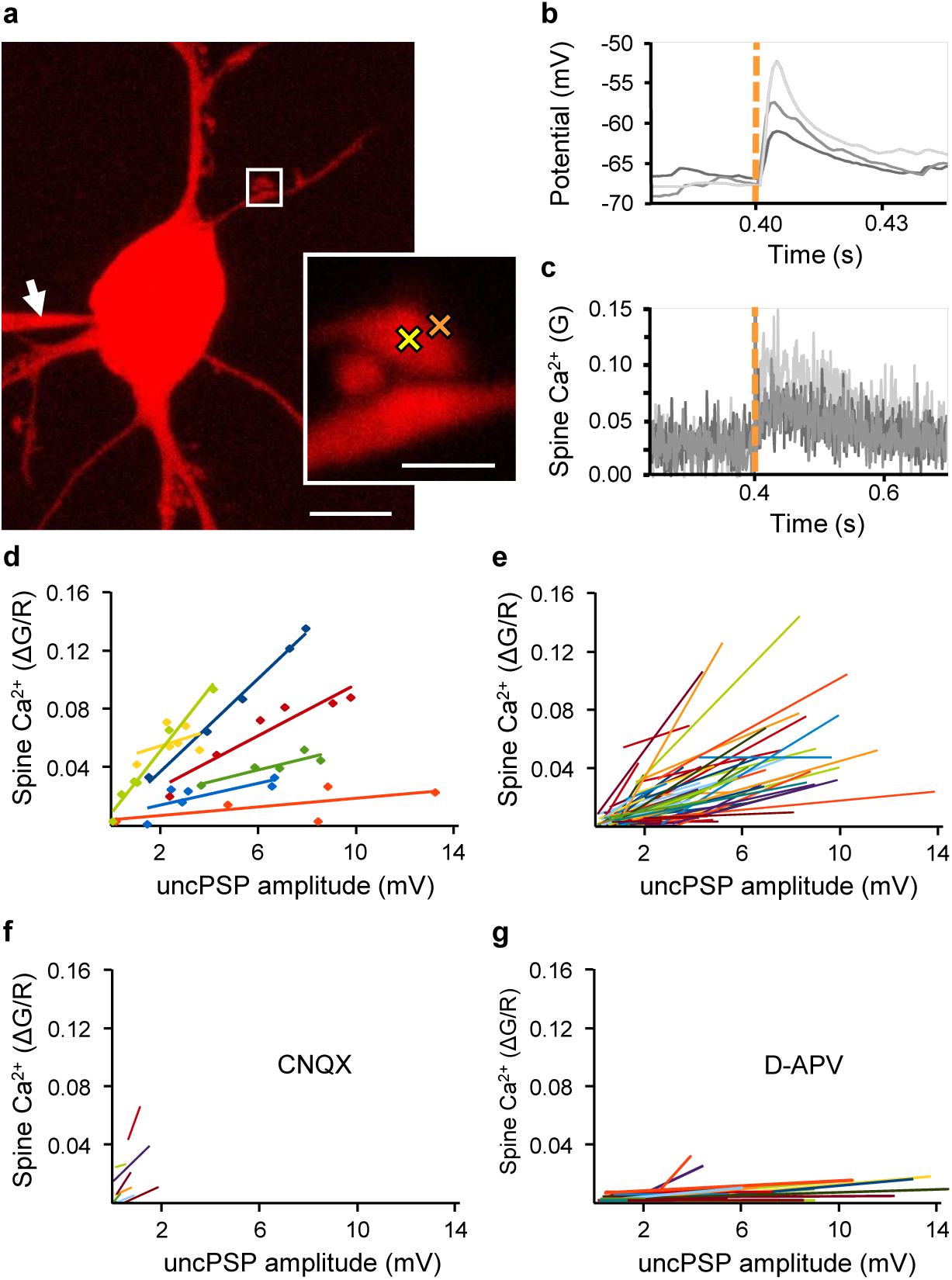
Heterogeneity of postsynaptic responses to glutamate uncaging. **a** Hilar mossy cell filled with Alexa 594 (red) and Fluo-5F for Ca^2+^ imaging (arrow points to patch pipette). Enlarged boxed area shows positions of imaging (yellow cross) and uncaging (orange cross) laser spots in the postsynaptic spine and the synaptic cleft, respectively. Glutamate was photo-released at the synaptic cleft from MNI-glutamate using two-photon uncaging. Scale bars: 20 μm and 5 μm, respectively. **b**, **c** Each synapse was probed with six laser pulses of increasing intensities, resulting in increasing amplitudes of uncPSPs and Ca^2+^ transients in the spine (responses to three laser pulses are shown). **d** Voltage and Ca^2+^ responses to increasing intensities (dots). Lines represent regression of peak Ca^2+^ amplitudes on uncPSP amplitudes in selected synapses (different colours). **e** Regression lines of all 79 synapses studied (in 79 slice cultures). **f** Application of the AMPA receptor antagonist CNQX diminished voltage responses (n = 8). **g** Application of the NMDA receptor antagonist D-APV drastically reduced the amplitudes of Ca^2+^ transients (n = 23).

Differences in synaptic states between individual MF synapses on MCs might be due to differences between slice cultures or MCs. Therefore, we early on made attempts to stimulate and record two MF synapses on the same MC. One of these experiments (n = 3) is shown in Fig. 1d-f. When we stimulated two different MF boutons contacting different spines on the same MC dendrite we noticed great differences in unitary EPSPs and postsynaptic Ca^2+^ transients between the two synapses on the same neuron. While stimulation of the synapse on spine 1 resulted only in subthreshold EPSPs, accompanied by Ca^2+^ transients of low amplitude, stimulation of the synapse on spine 2 led to frequent postsynaptic APs and large Ca^2+^ transients in the spine. Similarly, paired-pulse facilitation was different between the two synapses. These differences between the two synapses imply that the granule cell input to spine 2, but not that to spine 1, would *in vivo* be conveyed by the MC axon to large dorso-ventral portions of the ipsi- and contralateral dentate gyrus. We accordingly classified postsynaptic responses to bouton stimulation as subthreshold and suprathreshold, respectively, taking the different impact of a detonating synapse and a subthreshold synapse on network activity into account. MF synapses were thus of three response categories following the first stimulus: only subthreshold EPSPs, mixed sub- and suprathreshold responses, and only suprathreshold EPSPs (Fig. 2a, b, Supplementary Fig. 3). The fraction of suprathreshold responses of each synapse was positively correlated with the median EPSP slope but neither of these parameters was correlated with the dendritic distance of the synapse from the MC soma, indicating that the detonator status does not merely reflect proximity to the cell body (Supplementary Figs. 3 and 4a-h).

**Fig. 4.**
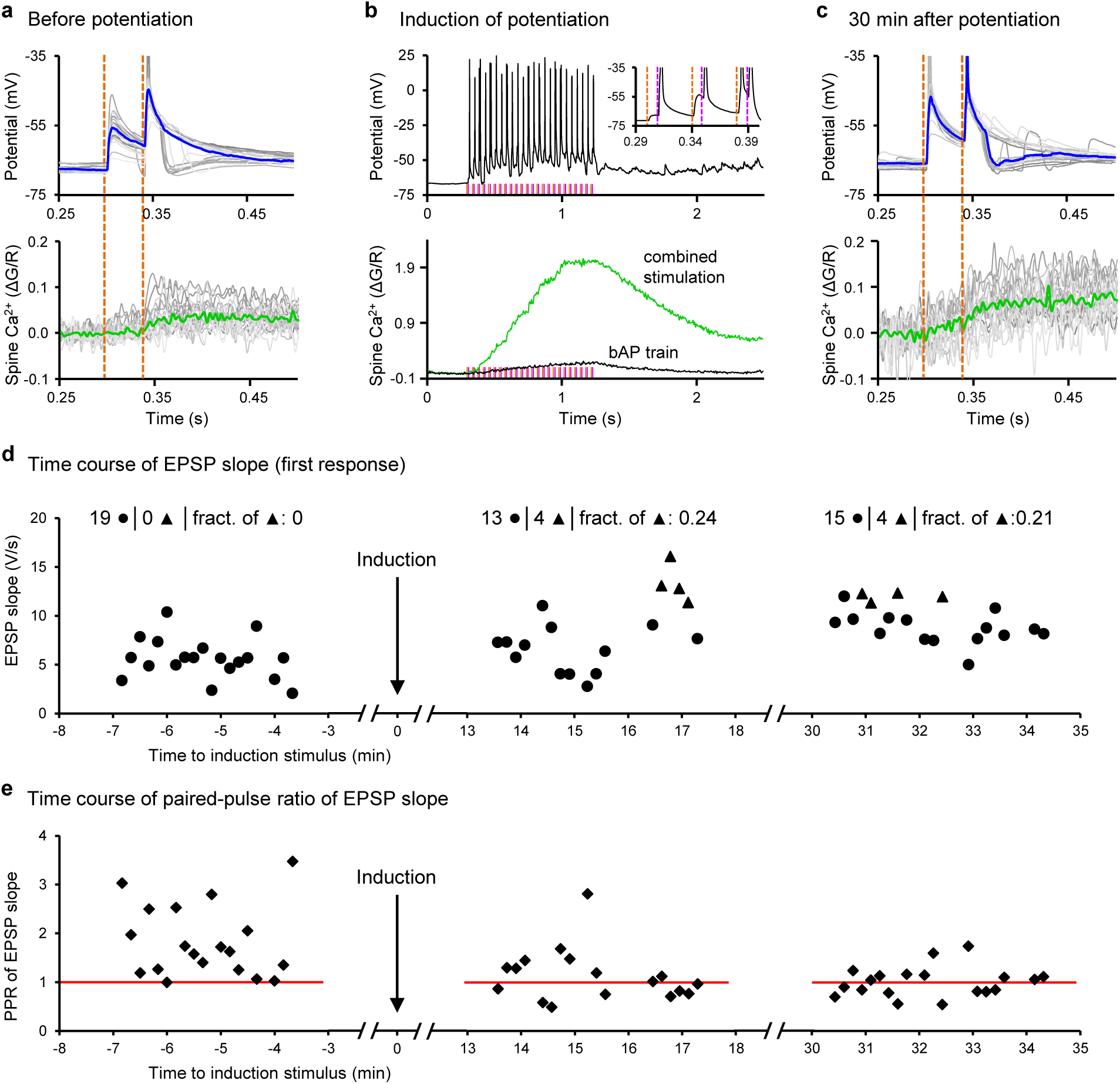
Potentiation of individual identified MF-MC synapses. **a** Unitary voltage (top) and Ca^2+^ response (bottom) to paired-pulse stimulation of a synapse prior to potentiation (synapse no. 3; blue and green lines: Medians of all repetitions). **b** Induction of plasticity in the synapse in **a** by combining bouton stimulation and backpropagating action potentials (bAPs); inset: first three responses. Orange dotted lines: Bouton stimulations; magenta lines: Postsynaptic stimuli (bAPs). Dramatic increase in Ca^2+^ during combined stimulation (green line) but not in response to a train of only bAPs (black line). **c** Same synapse as in **a**, 30 min after combined stimulation. **d** Time course relative to the induction train. EPSP slope in response to the first of paired-pulse stimuli, (●) subthreshold responses, (▲) suprathreshold responses; headlines: Numbers of sub-and suprathreshold responses, fraction of suprathreshold responses. **e** Time course of paired-pulse ratio of EPSP slope of the responses in **d**; red lines indicate a paired-pulse ratio of 1.

Responses to the first stimulus revealed a great variability of the EPSP slope and peak Ca^2+^ amplitude in the total of 26 synapses studied in 26 slice cultures, each colour-coded individually allowing for their identification in the various figures (Fig. 2b, Supplementary Fig. 3). Despite the observed variability, EPSP slope and peak Ca^2+^ response were positively correlated within single synapses (statistically significant correlation in 22 of 26 synapses, Supplementary Fig. 5) and among synapses (Fig. 2b). Of note, the different synaptic weights in response to repeated paired-pulse stimulations of a synapse were not correlated with the sequential order of repetitions, precluding accumulation of synaptic facilitation (no significant correlation of sequential order with EPSP slopes in 23 of 26 synapses and with peak Ca^2+^ in 21 of 26 synapses, Supplementary Fig. 5). The regression lines of EPSP slopes and peak Ca^2+^ responses differed remarkably between individual synapses, as did their amplitudes (Fig. 2a, b, Supplementary Fig. 3), pointing to variable glutamate release and variable AMPA and NMDA glutamate receptor contributions. Consistent with this hypothesis, varying the amount of glutamate in two-photon uncaging experiments resulted in corresponding changes in voltage responses (uncPSPs) and Ca^2+^ responses (Fig. 3a-c). Comparable to single bouton stimulation, we observed a great heterogeneity of the responses from all 79 synapses studied in these uncaging experiments (Fig. 3d, e). Application of the AMPA receptor antagonist CNQX almost completely abolished the uncPSP amplitudes, whereas treatment with the NMDA receptor blocker D-APV resulted in a drastic reduction of peak Ca^2+^ responses (Fig. 3f, g), indicating that EPSP amplitudes represent AMPA receptor activation and Ca^2+^ transients NMDA receptor currents at MF synapses similar to other synapses^42–44^. In contrast to single bouton stimulation, we did not observe suprathreshold responses to glutamate uncaging, reflecting that glutamate uncaging involved only a subset of postsynaptic densities contacted by a single MF terminal.

**Fig. 5.**
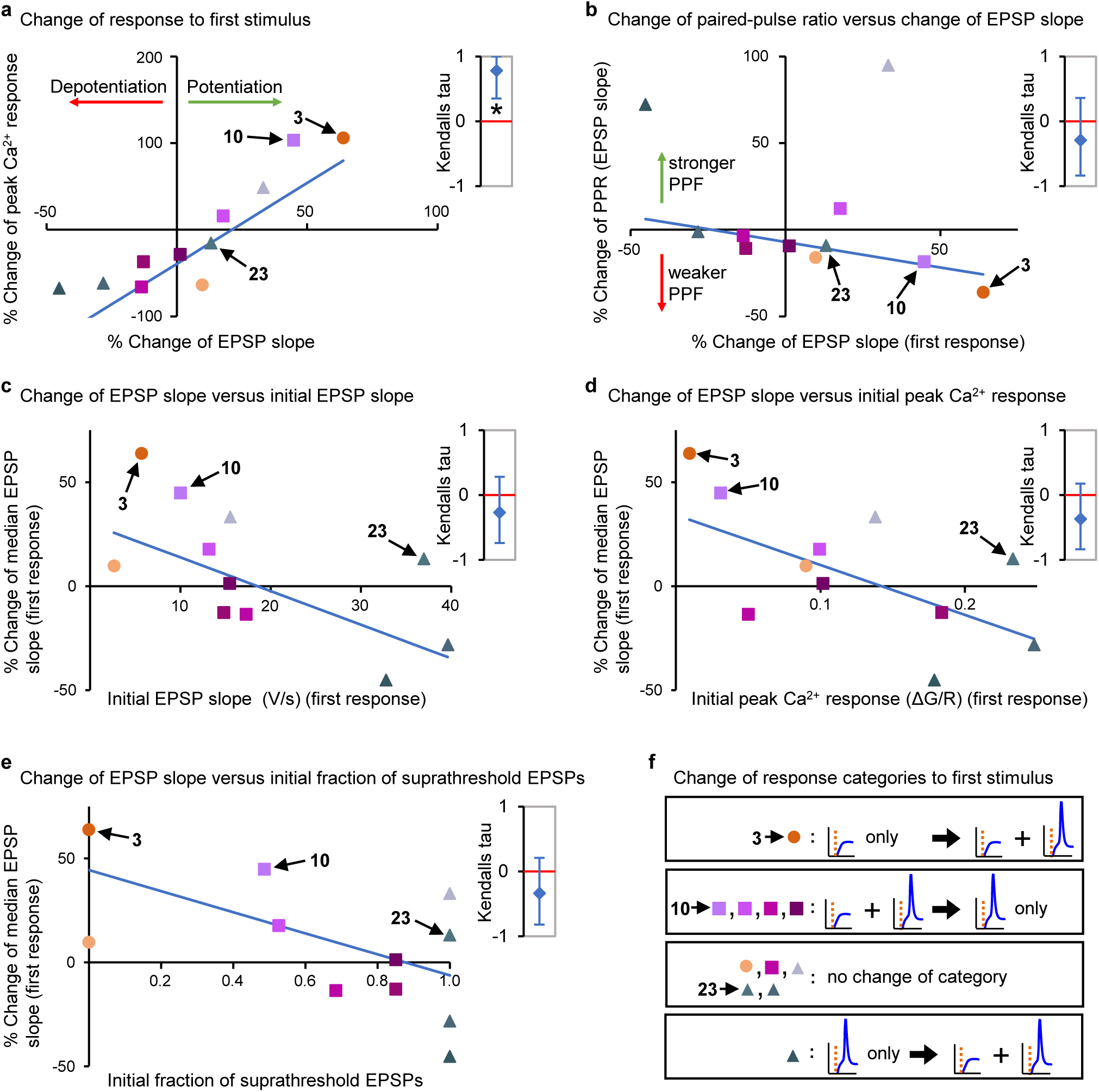
Heterogeneous synaptic plasticity of MF-MC synapses. **a** Percent change of slope of EPSP and peak Ca^2+^ amplitude in response to the first stimulus 30 min after potentiation in a subset of synapses; coding as in Fig. 2 (indicating the synaptic state prior to potentiation; medians of all repetitions per synapse). The synapses no. 3, 10, and 23 are marked as in Fig. 2. Blue line represents regression of percent change of peak Ca^2+^ on percent change of EPSP slope. Right panel shows the correlation between these parameters. The bootstrap mean (♦) of Kendalls tau and a confidence interval (0.025 – 0.975, vertical bars) is shown as a robust measure of correlation. If 0 (red line) is not included in the confidence interval, the parameters are considered to be correlated (*) at a significance level of 0.05. **b** Percent change of paired-pulse ratio of EPSP slope versus change of EPSP slope 30 min after potentiation of these synapses. Blue line represents regression, right panel shows the correlation between these parameters; PPF: Paired-pulse facilitation. **c** Percent change of EPSP slope 30 min after potentiation versus initial EPSP slope of these synapses. Blue line represents regression, right panel shows the correlation between these parameters. **d** Percent change of EPSP slope 30 min after potentiation versus initial peak Ca^2+^ response of these synapses. Blue line represents regression, right panel shows the correlation between these parameters. **e** Percent change of EPSP slope 30 min after potentiation versus initial fraction of suprathreshold EPSPs of these synapses. Blue line represents regression, right panel shows the correlation between these parameters. **f** Change of response categories of these synapses.

Presynaptic facilitation assessed by the paired-pulse ratio of EPSP slope revealed a contribution of the bouton to synaptic heterogeneity showing a strong facilitation of primarily weak synapses and weak facilitation or depression of strong synapses. Weak responses to the first stimulus were correlated with strong paired-pulse ratios of EPSP slope among synapses as well as within individual synapses, indicating that the variability of the first synaptic response is due to a variable initial release probability (Fig. 2c, Supplementary Figs. 3 and 6). Suprathreshold EPSPs following both stimuli occurred on average in 57% of all stimulations per synapse, whereas only one or no AP at all was observed in 23% and 19%, respectively (Table 1). Taken together, single bouton stimulation disclosed a great deal of heterogeneity of EPSP slopes and peak Ca^2+^ responses, pointing to individual contributions of single MF synapses to MC firing ranging from consistently subthreshold activation to robust detonation.

### Divergent plasticity following a uniform induction protocol

What determined the observed heterogeneity of MF synapses? The response to the first of paired stimuli represents the synaptic state at this particular point in time. Different strengths of synapses may indicate different histories of potentiation and depression. This would imply that the state of an individual MF synapse is the result of previous activity and is further modified by ongoing activity. Therefore, following the paired-pulse protocol, we induced long-term synaptic plasticity at these same synapses by a single train of 24 bouton stimulations (25 Hz), each paired with a delayed (+ 10 ms) backpropagating action potential (bAP) elicited in the postsynaptic cell (spike-timing-dependent plasticity^38^) (Fig. 4a-e). Our analysis of responses to paired-pulse stimulation, reporting changes in release probability and changes in EPSP slope and Ca^2+^ transients, allowed us to determine both presynaptic and postsynaptic plasticity (Fig. 5, Supplementary Fig. 7, Table 2). We noticed great differences in the changes of synaptic strengths and of paired pulse ratios 30 minutes after the combined stimulation (Fig 5a, b, Supplementary Fig. 7). Previously weak synapses showing initially small EPSP slopes, low peak Ca^2+^ amplitudes, and none or only a small fraction of suprathreshold EPSPs in response to the first of paired stimuli became potentiated, whereas previously strong synapses remained strong or became even depotentiated (Fig. 5c-e, Supplementary Fig. 7). However, all but one synapse showed an increase or no change in the fraction of suprathreshold EPSPs in response to the first of paired stimuli: Potentiation of 5 out of 11 synapses resulted in a change of the detonator status to the mixed sub- and suprathreshold and the all suprathreshold response category, respectively (Fig. 5f, Supplementary Fig. 7). Considering both responses to paired-pulse stimulation, the fraction of suprathreshold-only responses to both stimuli had increased significantly 30 min after induction of plasticity, whereas the fraction of paired-pulse responses containing at least one subthreshold EPSP decreased accordingly (Table 2). The paired-pulse ratio of EPSP slope decreased in most of the potentiated synapses. The synapses with the largest increase in EPSP slope showed the strongest reduction in paired-pulse ratio indicating that potentiation of these synapses was due to an increase in presynaptic release probability (Fig. 5b, Supplementary Fig. 7). Overall, these findings indicate that the outcome of plasticity depended on the initially encountered synaptic state of these MF synapses. Vice versa, our findings suggest that the initially encountered synaptic state had resulted from the different previous activities of these individual MF synapses.

**Table 2.**
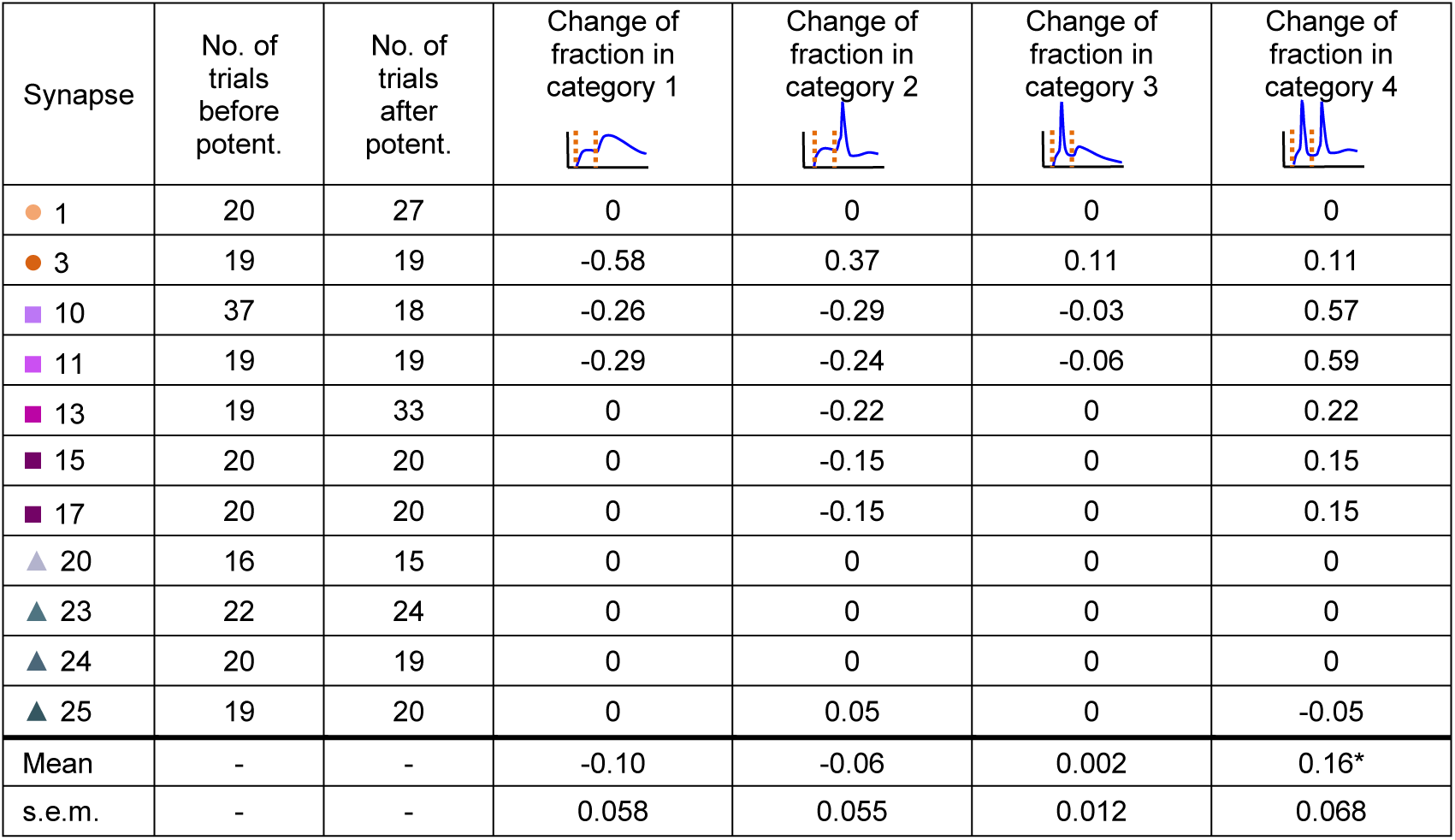
Change of response categories following potentiation. Response patterns are shown schematically on top (categories as in Table 1). Each synapse is represented in all 4 response categories (changes of fractions summing up to zero). Coding of synapses by coloured symbols as in Fig. 2. Bottom: Mean ± s.e.m. of fraction changes of the 11 synapses analysed. (*) significant increase in category 4 after potentiation (Wilcoxon signed-rank test with Pratts modification, n = 11, P = 0.03125).

| Synapse | No. of trials before potent. | No. of trials after potent. | Change of fraction in category 1 | Change of fraction in category 2 | Change of fraction in category 3 | Change of fraction in category 4 |
| --- | --- | --- | --- | --- | --- | --- |
| ● 1 | 20 | 27 | 0 | 0 | 0 | 0 |
| ● 3 | 19 | 19 | -0.58 | 0.37 | 0.11 | 0.11 |
| ■ 10 | 37 | 18 | -0.26 | -0.29 | -0.03 | 0.57 |
| ■ 11 | 19 | 19 | -0.29 | -0.24 | -0.06 | 0.59 |
| ■ 13 | 19 | 33 | 0 | -0.22 | 0 | 0.22 |
| ■ 15 | 20 | 20 | 0 | -0.15 | 0 | 0.15 |
| ■ 17 | 20 | 20 | 0 | -0.15 | 0 | 0.15 |
| ▲ 20 | 16 | 15 | 0 | 0 | 0 | 0 |
| ▲ 23 | 22 | 24 | 0 | 0 | 0 | 0 |
| ▲ 24 | 20 | 19 | 0 | 0 | 0 | 0 |
| ▲ 25 | 19 | 20 | 0 | 0.05 | 0 | -0.05 |
| Mean | - | - | -0.10 | -0.06 | 0.002 | 0.16* |
| s.e.m. | - | - | 0.058 | 0.055 | 0.012 | 0.068 |

## Discussion

In the present study, we provide evidence for a dynamic control of MC firing by single MF inputs that are separately regulated by state-dependent individual synaptic plasticity. Synaptic potentiation using an associative protocol was found to change synaptic strengths differentially, depending on the initially encountered state of the synapse. We probed the initial synaptic weight by monitoring the first response to paired-pulse stimulation and compared the response to the first stimulus before and 30 minutes after induction of synaptic plasticity. While this approach did not allow us to trace the history of the synaptic state back to before initial stimulation, we were able to follow the plastic response to a defined stimulus train, which was different in the synapses studied leading to potentiation but also depression. This bidirectional plasticity was observed pre-and postsynaptically and is consistent with the multitude of plasticity mechanisms reported for MF synapses^36,45–48^.

We simultaneously measured peak Ca^2+^ amplitudes and EPSP slopes in response to single MF bouton stimulations. These two parameters were correlated in individual synapses, but each synapse displayed a distinct relation between Ca^2+^ and unitary voltage responses (Fig. 2a, b, Supplementary Fig. 3). A likely explanation is that some MF synapses on MCs have many NMDA receptors while others have only very few. This is supported by glutamate-uncaging experiments, where we also found heterogeneity in Ca^2+^ to voltage relation, showing that this heterogeneity is due to postsynaptic mechanisms (Fig. 3d, e). Moreover, the uncaging experiments revealed that NMDA receptor blockade dramatically reduced Ca^2+^ transients, whereas blockade of AMPA receptors strongly decreased uncPSP amplitudes (Fig. 3f, g). Since glutamate uncaging was restricted to single spine heads, the heterogeneity in the ratio of peak Ca^2+^ amplitudes to EPSP slopes of synaptic responses could not be attributed merely to a potentially varying number of spine heads activated by a single MF bouton. Within single synapses EPSP slope was negatively correlated with paired-pulse ratio indicating that the observed heterogeneity of responses to the first pulse was based on a strong variability of release probability within single boutons (Supplementary Figs. 3 and 6). Between synapses we also found a pronounced heterogeneity of release probability as indicated by the paired-pulse ratios of EPSP slopes ranging from strong facilitation to depression. Thus, we encountered also the MF boutons in varying states corresponding to the postsynaptic heterogeneity (Fig. 2c).

**Fig. 6.**
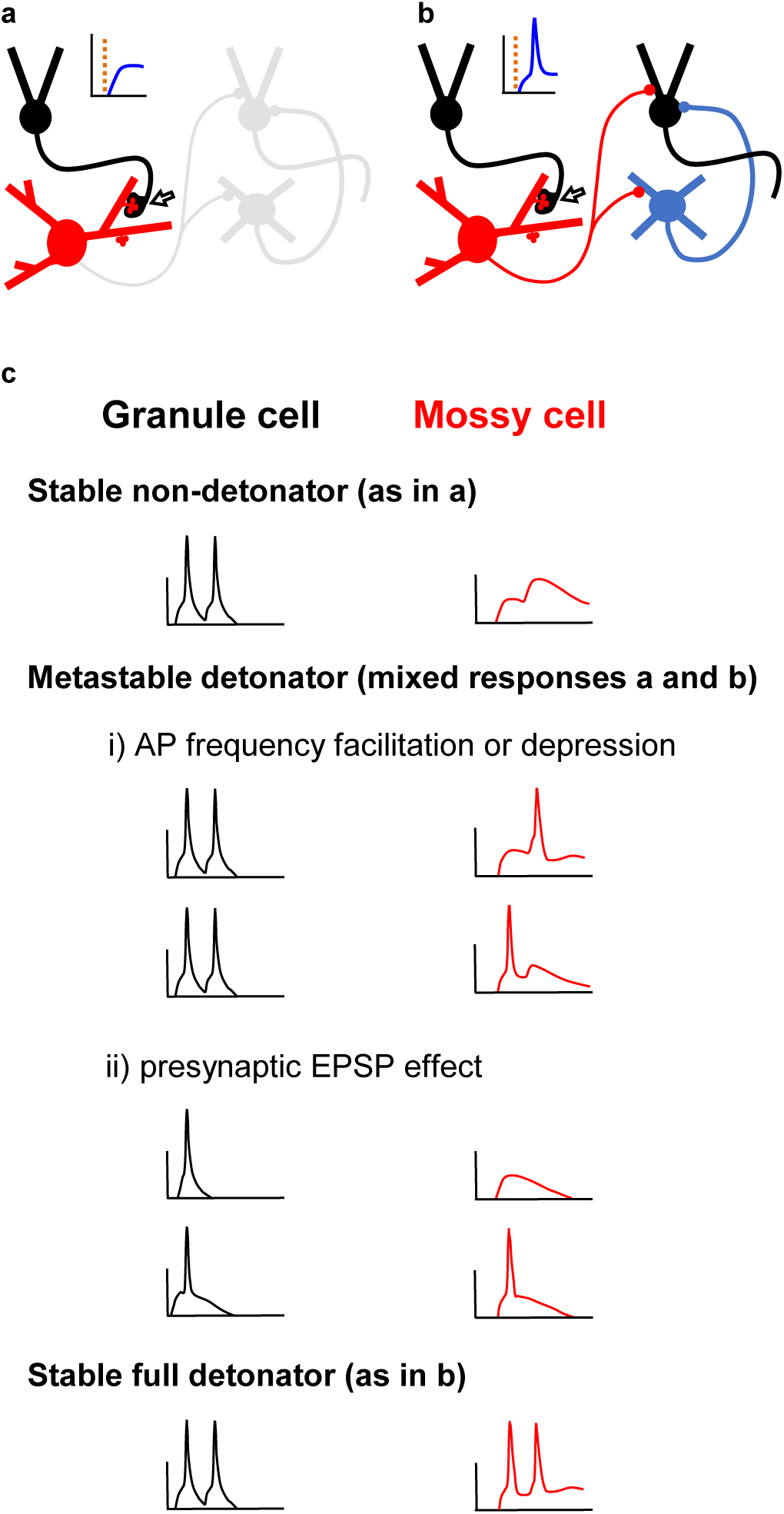
MF-MC synapses represent a metastable network switch. **a** If the synapse (open arrow) is in subthreshold mode, only the MC (red) will notice AP firing of the presynaptic granule cell (black). **b** In suprathreshold, detonating mode, granule cells and interneurons (blue) postsynaptic to the MC will be notified of this unitary MF-MC EPSP. **c** MF-MC synapses are encountered in three different states with respect to MC detonation. Stable non-detonator synapses show exclusively subthreshold responses. Metastable detonator synapses are on the edge to detonation. They may become facilitated or depressed and are dependent on AP frequency and accompanying presynaptic EPSPs^56^. Stable full detonator synapses consistently elicit AP firing of the MC (as illustrated in **b**).

Induction of plasticity at these synapses using an associative protocol resulted in correlated changes in peak Ca^2+^ amplitudes and EPSP slopes and corresponding changes in paired-pulse ratios even though some synapses were strengthened and others became weakened, pointing to a change in presynaptic glutamate release probability (Fig. 5). However, in 5 out of 11 synapses we observed pronounced changes in the Ca^2+^ to voltage relation, suggesting additional postsynaptic modifications (Supplementary Fig. 7). The direction of plasticity was not random, but depended on the initial state of the synapse, since weak synapses tended to be potentiated whereas already strong synapses showed depotentiation (Fig. 5c-e).

Plasticity at MF synapses has been shown to be expressed presynaptically, modulating glutamate release probability and resulting in potentiation^36^ or depotentiation^49^. Presynaptic GABAergic^50^, cholinergic^51^ and glutamatergic mechanisms are likely to contribute to MF bouton plasticity. More recent studies have revealed additional postsynaptic forms of plasticity at MF synapses, including NMDA receptor-dependent recruitment of NMDA receptors^46,52^, NMDA receptor-dependent potentiation of AMPA excitatory postsynaptic currents^47^, and NMDA receptor-dependent bidirectional spike-timing-dependent plasticity^53^. Depression of postsynaptic NMDA currents may result from the co-release of zinc from strongly activated MF boutons^54^. Given the heterogeneity in glutamate receptor content of MF-MC synapses reported here, one would expect heterogeneous responses to a defined stimulus, as was indeed observed when applying a defined plasticity protocol. Different forms of synaptic plasticity are generally attributed to different stimulation protocols. In MF-MC synapses we found that pre-and postsynaptic plasticity depended on the initial state of the synapse. At least for MF synapses on MCs, we have shown that there is a great diversity of synaptic states. Apart from the detonation property, which is a binary response variable, we found a continuum of synaptic states rather than discrete modes of transmission as suggested for synapses between CA3 pyramidal cells^55^. We observed plasticity of voltage and Ca^2+^ responses following a uniform protocol, spanning a wide range of changes without obvious clustering of response patterns, thus not supporting a shift between discrete modes of MF-MC synapses.

We conclude that the different behaviour of individual MF synapses is based on their individual functional history, but other factors may also contribute. The dendritic distance between the synapses and the soma varied to some extent although neither the fraction of suprathreshold responses nor the median slopes of the EPSPs did correlate with this distance (Supplementary Figs. 3 and 4). MF synapses are heterogeneous with respect to their structural components such as bouton size, the complexity of postsynaptic spines and the number of active zones^4,5^. However, increased structural complexity of MF synapses was found to result from their previous activity^24^. Thus, structural changes of individual synapses are likely to contribute to the observed functional heterogeneity of MF synapses on MCs.

We noticed different synaptic states with respect to postsynaptic MC firing in response to the first stimulus, ranging from subthreshold-only responses in one group of synapses to suprathreshold-only responses in another (Table 1, Fig. 2, Supplementary Fig. 3). While a suprathreshold MF synapse on a MC may eventually result in the activation of large populations of granule cells and interneurons in distant lamellae in the ipsi- and contralateral hippocampus, these same neurons will not be activated by a subthreshold MF-MC synapse (Fig. 6a, b). Fifteen out of the 26 synapses show both suprathreshold and subthreshold responses to the first of paired stimuli. One is tempted to speculate that these mixed synapses are toggling between sub- and suprathreshold responses due to the analogue control of release probability by the integration of presynaptic granule cell EPSPs in MF boutons^56^. Thus, MF-MC synapses might function as a metastable network switch, depending on the granule cell firing rate as well as the strength of a granule cell EPSP accompanying the presynaptic AP^56^ (Fig. 6c). In response to an associative induction protocol many synapses showed an increased fraction of suprathreshold EPSPs, suggesting that synaptic activity changes the detonator status of MF-MC synapses (Table 2, Fig. 5f, Supplementary Fig. 7). We conclude that MF-MC synapses integrate previous activity to become stronger or weaker, thus controlling modulation of the dentate network by switching between detonating and non-detonating MC activation. The results imply that single granule cells control MC firing in a precisely regulated manner reflecting their synaptic input patterns. Direct detonation of MCs by single granule cells as shown here is well compatible with previous reports on the high firing rate of MCs *in vivo* when compared to the sparse firing of granule cells^15,18,40^.

Detonation of a MC by a single granule cell promotes di-synaptic inhibition of dentate granule cells locally and excitation as well as di-synaptic inhibition of granule cells in distant parts of the dentate gyrus^8^. A single bursting granule cell having established a full detonator synapse with a MC may potentiate the MCs output synapses thereby shifting the excitation/inhibition balance of large numbers of distant granule cells postsynaptic to that MC^21^. Recent optogenetic studies provided evidence for MCs being the first excitatory input to adult-generated granule cells when their growing dendrites reach the inner molecular layer, the termination zone of distal MC axons in the ipsi- and contralateral dentate gyrus^23^. Thus, the MC input may be important for the network integration of newly generated granule cells, which have been shown to mediate pattern separation, whereas old granule cells facilitate pattern completion^57^. Remarkably, genetic ablation of MCs resulted in impaired pattern separation^22^, suggesting that MCs controlled by the MF input synapses play a crucial role in the integration of new granule cells and contextual discrimination mediated by these neurons. The conditional detonator synapses of MFs on CA3 pyramidal cells are known to be involved in complex network functions such as pattern completion and pattern separation^57–59^. The metastable direct detonator MF-MC synapses introduced here may accordingly play a critical role in the functional organization of the dentate gyrus.

## Methods

### Preparation of organotypic slice cultures

Mouse pups of both sexes (P4 to P5; C57bl6, local colony) were decapitated with scissors in accordance with institutional, European and international guidelines (licence no. ORG_582). The entorhino-hippocampal complex was dissected out of the brains and cut horizontally at 300 μm thickness using a Mc Illwain tissue chopper. Slices containing the entorhinal cortex attached to the ventral hippocampus were transferred to Millicell inserts (Millipore, Schwalbach, Germany)^60^ and incubated for at least 28 days at 37 °C and 5% CO_2_ in 1.2 ml of a medium composed of 50% Minimum Essential Medium, 25% Basal Medium Eagle, 25% heat-inactivated horse serum, 2 mM glutamine, and 6.5 mg/ml glucose with pH adjusted to 7.3. The medium was exchanged three times per week. Substances were obtained from Invitrogen (Darmstadt, Germany).

### Patch clamp recording solutions

Slice cultures were superfused at 35 °C (measured 1 mm beneath the slice) with artificial cerebrospinal fluid (ACSF) containing 125 mM NaCl, 25 mM NaHCO_3_, 2.5 mM KCl, 1.25 mM NaH_2_PO_4_, 1 mM MgCl_2_, 25 mM glucose, and 2 mM CaCl_2_, pH adjusted to 7.30 with NaOH, ~305 mOsm, equilibrated with 95% O_2_ and 5% CO_2_.

The MNI-glutamate uncaging was performed in a self-contained volume of 0.7 ml ACSF composed of 120 mM NaCl, 25 mM NaHCO_3_, 2.5 mM KCl, 1.25 mM NaH_2_PO_4_, 1 mM MgCl_2_, 10 mM glucose, and 2 mM CaCl_2_, and 20 mM 4-Methoxy-7-nitroindolinyl (MNI)–glutamate (Tocris, via BIOZOL, Eching, Germany). We needed a high concentration of MNI-glutamate for uncaging to keep the uncaging pulses short while minimizing laser intensities to mimic glutamate release kinetics without damaging the synapse. Moreover, multiquantal release at MF synapses^61^ requires uncaging of larger amounts of glutamate when compared to the small Schaffer collateral synapses in CA1. Glutamate receptor antagonists (200 μM 2-amino-5-phosphonopentanoic (D-APV) and 20 μM 6-cyano-7-nitroquinoxaline-2,3-dione (CNQX) (both from ascent scientific, Cambridge, U.K.) were present in some uncaging experiments. The solution was covered by a constant flow of moistened gas (95% O_2_ and 5% CO_2_) and was heated locally to ~35 °C. Evaporation was compensated for by continuous addition of water using a roller tube pump to keep osmolarity at ~ 305 mOsm.

The pipette solution for whole-cell recordings contained 133 mM K-gluconate, 12 mM KCl, 10 mM HEPES, 7 mM Na_2_-phosphocreatinine, 4 mM Mg-adenosine-5-triphosphate, 0.3 mM Na_2_-guanosine-5-triphosphate, 9 mM sucrose (to achieve ~ 305 mOsm), pH adjusted to 7.30 with KOH, and either 200 μM Fluo-5F and 100 μM Alexa-594 dextran (10 kD) for glutamate-uncaging experiments or 800 μM Fluo-4FF and 40 μM Alexa-594 dextran (10 kD) for the rest of the experiments. Electrode tips were filled with intracellular solution not containing the dyes in order to avoid excessive background labelling of the tissue before seal formation. We used Alexa 594 dextran rather than the hydrazide form for intracellular labelling to avoid effects of the carbonyl-reactive hydrazide^62^ and used different Ca^2+^ indicators with different affinities since we expected larger Ca^2+^ transients in the bouton stimulation experiments^63^.

The pipette solution for loose-seal MF bouton stimulation contained 145 mM NaCl, 2.5mM KCl, 1 mM MgCl_2_, 2 mM CaCl_2_, 10 mM HEPES, pH adjusted to 7.40 with NaOH, and 50 μM Alexa 488 hydrazide. Since dextran conjugates are actively taken up by neurons from external solutions, we used Alexa 488 hydrazide for transient labelling of the extracellular space for shadow patching. Albumin (Albumin bovine Fraction V, 25 mg/ml, SERVA, Heidelberg, Germany) was added to reduce seal resistance. Chemicals were obtained from Sigma-Aldrich (Taufkirchen, Germany) and dyes from Invitrogen.

### Electrophysiology

Mossy cells in the hilar region of the hippocampus were visualized under video IR-Dodt contrast^64^, and whole-cell access was established with 5-10 MΩ patch pipettes fabricated from borosilicate glass capillaries (outer diameter 2 mm, inner diameter 1 mm; Hilgenberg, Malsfeld, Germany) using a small piezo-driven micromanipulator (Kleindiek-nanotechnik, Reutlingen, Germany) for long-term stability of the recordings. On break in, resting membrane potentials ranged from −63 mV to −78 mV (−55 mV to −65 mV in MNI-glutamate solution). Cells were recorded in current clamp mode using an ELC-03X amplifier (npi electronic, Tamm, Germany) in bridge mode. Series resistance and pipette capacitance compensation were adjusted using a phase-sensitive technique^65^. Amplifier output was low-pass Bessel-filtered at 20 kHz and digitized at 100 kHz using a PC-board (NI PCI6251, National Instruments, Munich, Germany) and in-house software based on LabVIEW^TM^ (National Instruments). The input resistance of MCs was determined by the ratios of the steady state voltage responses to small current steps (200 ms) ranging from −20 to +20 pA. Membrane time constant was derived from exponential fits of the voltage response to hyperpolarizing current steps (150 ms, −20 pA). AP firing threshold was defined independently of synaptic stimulation as the mean membrane potential right before the first elicited APs in response to depolarizing current steps of increasing amplitude (iterated until five steps with AP responses were obtained). Back-propagating action potentials (bAPs) were elicited by short somatic current injections (1.5 nA, 1.5 ms).

Labelled spines of the patched mossy cells were approached using a second pipette (theta borosilicate glass, outer and inner diameter 2 mm and 1.4 mm, respectively; Hilgenberg) filled with Alexa 488 hydrazide in the extracellular solution. Presynaptic MF boutons were identified as shadows covering labelled spines, contrasting against the stained extracellular space^35^, and loose-seal cell-attached mode was established^66,67^. MF boutons were stimulated with 0.5 ms pulses using a second ELC-03X amplifier (npi electronic) in current clamp x 10 mode (median: 36.6 nA, range 15 to 75 nA). Adequate stimulation intensity was determined for each bouton by increasing the current amplitude until an EPSP was recorded reliably in the postsynaptic cell with a maximal delay of 1.0 ms following the end of the stimulus (mean delay: 0.546 ms ± 0.188 s.d.; 26 synapses). Voltage and Ca^2+^ responses were all-or-none with amplitudes independent of the bouton stimulation intensity (Supplementary Fig. 1a-c). Due to the large stimulus artefact recorded in the presynaptic pipette we were not able to monitor the AP elicited in the bouton in loose-seal cell-attached mode. Therefore, we did not attempt to determine failure rates using stimulation at threshold. During presynaptic stimulation a very small depolarization of the postsynaptic cell by less than 0.2 mV was recorded that typically returned back to the pre-stimulus potential before the onset of the EPSP, indicating synaptic activation rather than direct stimulation of the postsynaptic cell (Supplementary Fig. 1a-c). Moreover, contamination of the recorded EPSPs with synaptic responses following antidromic AP propagation to the same MC is unlikely, given the spacing of MF boutons^68^ and a propagation speed of ~ 210 μm/ms^69^. Indeed, we did not observe abrupt increases of the voltage slope during the rising phase of the EPSPs that would have been indicative of such compound signals.

Paired-pulse stimuli with an inter-stimulus interval of 40 ms were applied to test pre- as well as postsynaptic components of synaptic transmission. Paired-pulse stimulation was repeated at least 20 times (interval ≥ 10 s) to assess variability of synaptic transmission at a given synapse. This time interval proved to be sufficiently long to avoid unintended short-term potentiation of synapses during the repetitions (Supplementary Fig. 5). A single train (25 Hz) of 24 bouton stimuli, each combined with a delayed (+ 10 ms) bAP elicited in the postsynaptic cell, was applied to induce synaptic plasticity (Fig. 4b). This stimulation pattern resembles activity observed in granule cells *in vivo*^40,70^ and is known to involve postsynaptic mechanisms such as spike-timing-dependent plasticity^38^ at MF synapses^46,47,53^. 11 out of 26 synapses were successfully re-evaluated 30 min after the induction protocol applying again at least 20 paired-pulse stimuli. Some synapses were additionally tested 15 min after the induction (Fig. 4d, e). One synapse was analysed per slice culture with the exception of those experiments aimed at analysing different MF synapses contacting the same mossy cell.

### Imaging and glutamate uncaging

The recording chamber was integrated into a custom-made two-photon laser scanning microscope based on a BX50WI microscope frame and a Fluoview 300 scanning unit (Olympus, Hamburg, Germany). Excitation laser light was delivered through a 60 x water immersion objective lens (NA 1.1, LUMF, Olympus). The excitation laser beams were expanded to overfill the back pupil of the objective at least twofold to achieve maximal spatial resolution. Fluorescence was detected by photomultiplier tubes (PMT, R3896; Hamamatsu Photonics, Herrsching, Germany) in non-descanned mode through the objective lens (green light: dichroic 680 DCXXR, 2 mm AR coated BG39; Schott, Mainz, Germany, and band-pass HQ 510/80, AHF analysentechnik, Tübingen, Germany) and an oil immersion condenser (NA 1.4, BX-UCDB-2, Olympus; green and red light: IR blocker HC-2P Emitter 680/SP, dichroic 560 DCXR, band pass HQ 510/80 and HQ 610/75, all from AHF analysentechnik). The amplified PMT output signals were digitized at 2.5 MHz using a PC Board (NI PCI-6132, National Instruments). Scanning was controlled using Fluoview software (Olympus). Timing and data acquisition were accomplished using a program implemented in LabVIEW™ and the NI PCI-6251 board (National Instruments). The two-photon excitation source for imaging was a Ti:Sa laser tuned to 805 nm with a pulse repetition rate of 1 GHz and a spectral full width at half maximum of ~ 30 nm (Gigajet 20C; Laser Quantum, Konstanz, Germany; pump laser Verdi V5; Coherent, Dieburg, Germany; pulse length at focal plane ~ 1.2 ps). Laser light for uncaging (720 nm, 76 MHz pulse repetition rate, ~ 5 nm pulse width) was generated in a Mira 900F (Coherent; pump laser Verdi V10; pulse length at focal plane ~ 0.4 ps). Excitation laser beams were combined using polarizing optics. A Keplerian telescope comprised of two biconvex lenses of 400 mm focal length was used to locate steering mirrors for both laser paths in front of the combining beam splitter in a conjugate plane to the scanning mirrors and the back focal plane of the objective lens, respectively^71^. This allowed convenient repositioning of the uncaging spot with respect to the imaging spot. These imaging conditions improved the signal to noise ratio due to a higher tolerable excitation level. We used point scans rather than line scans to gather maximal time resolution and to avoid dead times during recordings. As a result, individual recordings were analysed without averaging. Laser power was controlled by means of a half-lambda wave plate in the imaging laser path and an electro-optical modulator (Conoptics, via Polytec, Waldbronn, Germany) in the uncaging laser path. The dark current of the PMTs was measured over 100 ms before opening of the shutter (VS-25; Vincent Associates, via Acal BFi, Gröbenzell, Germany). Baseline fluorescence was collected for 200 ms before electrical and 300 ms before optical (0.5 ms) stimulation. Fluorescence response to the stimuli was monitored for 1 s (paired-pulse bouton stimulation), 1.5 s (uncaging), and 2.2 s (combined stimulation protocol). The uncaging laser spot was moved over the circumference of a spine to find the synaptic contact (no response if no hit). Background fluorescence was measured in the unlabelled tissue close to the labelled spines before stimulation onset.

The dendritic distance was determined as dendritic path length between a synapse and the MC soma using z-projections of two-photon image stacks (voxel size: 0.184 × 0.184 × 1.0 μm; Supplementary Fig. 4a, b).

### Exclusion criteria for quantitative analysis

Traces were excluded from analysis if one of the stated criteria was not met: Spontaneous AP or spontaneous synaptic event at the recorded synapse not closer than 0.1 s before the first stimulus, to allow for the determination of basal Ca^2+^-related fluorescence; maximal tolerated baseline fluorescence (green by red): 1.0; maximal tolerated postsynaptic holding current: −0.5 nA; maximal tolerated postsynaptic membrane potential before first stimulus: −60.0 mV.

### Analysis of EPSPs

Voltage recordings were subjected to a smoothing procedure based on locally adaptive Bayesian P-splines (algorithm “FlexNEG”)^72^ in order to reduce noise effectively without flattening of steep changes. Onset of EPSPs was determined iteratively using derivatives of decreasingly smoothened copies of nested intervals of a short time period (1 – 2 ms) following the stimulus. The maximal slope of the rising EPSP (before AP firing in case of suprathreshold EPSPs) was determined similarly using decreasingly smoothened copies of nested intervals of the extracted rising phase. The threshold potential of synaptically evoked APs was defined as the inflection point of the rising phase of the AP, again obtained iteratively in nested intervals of a short time interval before AP peak.

### Quantification of fluorescence intensity recordings

Relative changes in fluorescence in response to stimulation were calculated as green fluorescence minus green baseline fluorescence divided by red fluorescence (∆G/R), which is approximately proportional to the elevation of cytoplasmic [Ca^2+^] over baseline^63^. Individual traces were smoothened by a 5 ms wide rectangular moving average filter. Traces were fitted to a sum of exponential peak functions^73^ using a genetic algorithm. Peak Ca^2+^ responses were determined separately for each stimulus as amplitudes of these peak functions.

### Statistics

The Theil-Sen estimator of the regression slope and accordingly of the regression intercept was used for robust regression of EPSP slopes on peak Ca^2+^ responses (number of samples for regression estimate = 10000). A confidence interval (0.025 – 0.975, equivalent to α = 0.05) for Kendall’s tau was obtained using the percentile bootstrap method (number of bootstrap samples = 1000) as a robust estimate of correlation of parameters between synapses as well as within synapses, respectively^74^. Changes of fractions within categories of synaptic responses on paired-pulse stimulation following potentiation were tested under the null-hypothesis that there is no change using Wilcoxon signed-rank test with Pratts modification for zero differences (α = 0.05; function “wilcoxsign_test (x ~ y, distribution = "exact", zero.method = c("Pratt")” from R-package “coin”^75^, called via R-script^76^). All procedures were implemented in LabVIEW™ (National Instruments).

## Data availability

The data that support the findings of this study are available from the corresponding author upon request.

### Acknowledgements

We thank Prof. Peter Jonas, Prof. Thomas Oertner and Prof. Roger Nicoll for critically reading an earlier version of the manuscript, Prof. Imre Vida for helpful discussion, and Bettina Herde, Janice Graw and Dung Ludwig for excellent lab assistance. Supported by the Deutsche Forschungsgemeinschaft (FOR 2419: FR 620/14-1) and Landesforschungsförderung Hamburg (LFF-FV27b) to M.F.; M.F. was Research Professor for Neuroscience of the Hertie Foundation.

## Author contributions

A.D. designed the experiments, developed targeted “shadow” patching of MF boutons together with U.M., and performed all analyses and programming. U.M. performed the experiments combining bouton stimulation, mossy cell recording and two-photon Ca^2+^ imaging and contributed to experiment planning. A.T. performed the glutamate uncaging experiments. M.F. together with A.D. developed the concept of the paper and wrote the manuscript.

